# Towards universal models for collective interactions in biomolecular condensates

**DOI:** 10.1101/2024.10.17.618883

**Authors:** Edoardo Milanetti, Karan KH Manjunatha, Giancarlo Ruocco, Amos Maritan, Monika Fuxreiter

## Abstract

A wide-range of higher-order structures, including dense, liquid-like assemblies, serve as key components of cellular matter. The molecular language of how protein sequences encode the formation and biophysical properties of biomolecular condensates, however, is not completely understood. Recent notion on the scale invariance of the cluster sizes below the critical concentration for phase separation suggests a universal mechanism, which can operate from oligomers to non-stoichiometric assemblies. Here we propose a model for collective interactions in condensates, based on context-dependent, variable interactions. We provide the mathematical formalism, which is capable describing growing dynamic clusters as well as changes in their material properties. Furthermore, we discuss the consequences of the model to maximise sensitivity to the environmental signals and to increase correlation lengths.

## I. INTRODUCTION

Our understanding of living matter has been revolutionised by the notion on higher-order protein structures^1–3^ (Figure 1). These nonstoichiometric assemblies regulate key cellular processes and can sample a wide-range of dynamics and material states^4^. Changes in particular in their biophysical properties may shift the physiological to an aberrant state, leading to dysregulation, as implicated in numerous human diseases^5,6^. The conversion of liquid-like to solid-like assemblies, for example to amyloid aggregates, is a hallmark in neurological disorders, such as Alzheimer’s or Parkinson’s disease or amyotrophic lateral sclerosis (ALS)^7,8^. The contribution of dense, liquid-like clusters, often termed as biomolecular condensates, to physiological processes is increasingly recognised^9,10^. Nevertheless, our understanding of the molecular nature and interactions of condensates is incomplete. Growing experimental evidence also highlight that these clusters also can be formed below the critical concentration of liquid-liquid phase separation^11–13^. In this regard, it is important to emphasize that most of the concepts related to phase separation and demixing (or liquid-liquid phase separation) are developed, in the vast majority of cases, for binary systems. In contrast, in cellular environments, the system is composed of a large number of different components, each of which is present in small quantities. In the formation of droplets of liquid *A* within the context of liquid *B* the fluctuation in the number of molecular components plays a fundamental role. Since these components are present in small numbers of molecules, they may follow a non-Gaussian distribution, and in certain extreme cases, may be limited to one or even no molecules. This leads to variations in the properties of liquid *A* from droplet to droplet, which tends to *blur* the phase separation boundaries in the phase diagram.

**FIG. 1.**
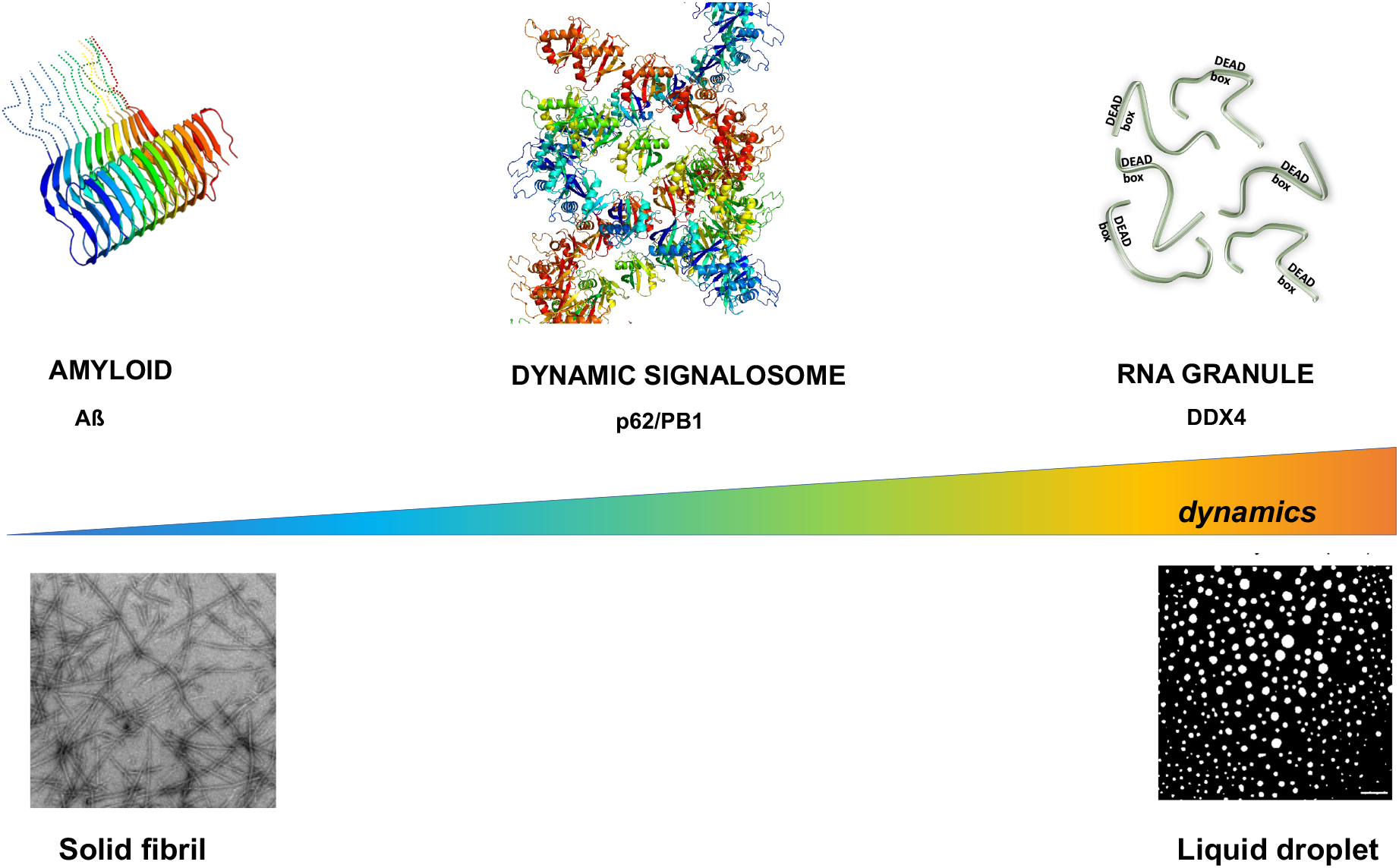
Higher-order protein structures sample a wide-range if dynamics and material states self-organization of which is stabilized by collective interaction. In this article we argue that fuzzy binding mechanisms can generate such collective interaction network and describe the main characteristics of these higher-order states. The hierarchy of self-organization is shown along increasing dynamics and interaction heterogeneity (*from left to right*): amyloid (Amyloid *β* ; PDB: 2mxu), dynamic signalosome (p62 PB1 domain; PDB: 4uf9), liquid-like condensate (DDX4 nuage granule). The conversion between liquid droplets and hydrogels (not shown) is reversible, while transition to amyloids is and irreversible process. Formation of Intermediate states, like functional aggregates or dynamical signalosomes is also reversible.

There are fundamental open questions regarding the organisation of condensates: *i)* whether interactions driving condensate formation are different from those, which assemble stoichiometric protein complexes? *ii)* What is the molecular basis of their *collective* nature? *iii)* How the selectivity of condensate interactions is controlled? *iv)* What is the basis of their cellular context-dependent regulation, assembly/disassembly? *v)* Which are the consequences of the condensate interactions in terms of macroscopic behavior and physical properties? *vi)* what is the effect of the large concentration fluctuation from droplet to droplet?

Numerous studies have attempted to delineate the interactions distinguished in formation of condensates and identify the key motifs, which drive this process^14^. A wide variety of interactions, including electrostatic^15^ and van der Waals interactions^16^ as well as *π*− *π* contacts^17,18^ were found to be critical for droplet formation. Our motivation in this article is different. We focus on the global, physical features of the condensate interaction network, and aim to provide general models, which can account for these characteristics. We aim at understanding: *i)* the dynamic nature of the interactions as droplets fuse/fission; *ii)* the dependence of droplet formation/disassembly on the external factors; *iii)* the possible change in biophysical properties of the droplets. Here we aim to define the boundary conditions for interactions, which enable these emergent properties and deduce the phenomenological consequences of the corresponding interaction network.

Our starting point is the recent analysis of experimental data of various proteins, which were observed to form dense, sizeable droplets below the critical concentration for phase separation^11–13^. The droplet size distributions could be described by a scale-invariant log-normal function, which could also estimate the critical concentration for protein phase separation^19^. The existence of scale invariance for the droplet size distribution, suggests a universal behaviour^20^, independent of the sequences and structures of the proteins undergoing phase separation. This is in line with the earlier notion on the generic nature of protein condensates^21^. With respect to the interactions, scale invariance stems from self-similarity at different length scales^22^, leading to universality.

The article will be organised as follows. First, we describe protein interactions at different scales (Section II). Second, we discuss the organisation of collective, dynamic interactions (Section III). Third, we analyze the sensitivity of the interaction network to environmental conditions (Section IV). Fourth, we detail the phenomenological consequences such as the increased correlation length and scale-invariance (Section V). Appendix A describes the mathematical formalism for collective interactions; and Appendix B details the formalism for scale invariance and correlation length. Finally, we conclude the article with the possible applications of our model in particular for therapeutic approaches and drug discovery.

### II. BRIDGING BETWEEN DIFFERENT SCALES OF PROTEIN INTERACTIONS

Protein interactions appear to be distinct at small and large scales of molecular organisation. Stoichiometric protein complexes are held together by well-defined contacts between specific residues, while biomolecular condensates are assembled through multiple valances, with a wide diversity of chemical nature and affinities. Which interactions can operate at both scales?

We initiate by asking how the external conditions (laboratory or cell) influence the activity of stoichiometric protein complexes. Fine-tuning of biological activities in accord with diverse signals often involves spatiotemporal variations at the interface formed by specific partners^23,24^. Cell cycle regulation by protein phosphatases, for example, is associated with dynamic, transient interactions by the regulatory loop, which can simultaneously bind different inhibitors^25^. In a similar vein, transcriptional co-activators may simultaneously engage with different activator domains through heterogeneous interactions by the same target site^26,27^. Intriguingly, transcription activity reduces upon increasing stability of the binding motifs, which reduces heterogeneity of the bound complex^3^. Although seems counterintuitive, advanced structure determination and biophysical techniques demonstrate that specific association can be achieved through multiple conformations^28^. For example, through multiple orientations of a short secondary structure, which is accommodating into a shallow binding cleft^29^. Although each orientation generates a distinct contact pattern, the bound ensemble involves a specific set of residues, which interact in a combinatorial manner. Accumulation of similar residues, which compete for the same target results in ambiguous, sub-optimal bound states, for example, in case of consecutive negative residues bind to a single positive residue of the partner^30^. Indeed, interaction fuzziness is analogous to the rugged energy landscape of protein folding, which is detailed elsewhere^31^.

Fuzzy interactions are enabled by evolutionary conserved, simple biochemical features, while the amino acid types are only weakly constrained^27,32^. An evolutionary selection for fuzzy binding was proposed in complex processes, which simultaneously affect multiple pathways^33^. *Multivalency* appears to be consistent and pertinent feature of proteins^34^, which form condensates, although the underlying binding motifs (i.e. *valence*) may not be recognised^14^. Simple features, for example patterning of residues with distinct physico-chemical characters can be sufficient to observe liquid-like droplets^35^. Although high resolution structural characterisation of condensates is still lacking, conformational heterogeneity of the assembly is supported by various lines of experimental evidence (eg. solution and single molecule methods)^36,37^. Conformational heterogeneity can be generated by combinatorial binding mechanisms via multiple valances^38^ as in case of prion-like domains^39^, or via other fuzzy binding mechanisms (eg. multiple orientations using the same site) by more structured proteins^40,41^. In both cases, high binding entropy can be attained in line with the dynamics and often liquid-like character of the condensates^21^.

Taken together, variable, fuzzy interactions can generate self-similarity at different length-scales of protein organisation and may serve as a universal feature.

## III. ON THE NATURE OF COLLECTIVE, DYNAMIC INTERACTIONS

In this section we address the problem of dynamic networking within the clusters formed below the critical concentration of phase separation. The distribution of droplet sizes of various proteins, including Fus^11^, alpha-synuclein^12^ and the nuage granule protein DDx4^42^ can be described by a scale invariant distribution as

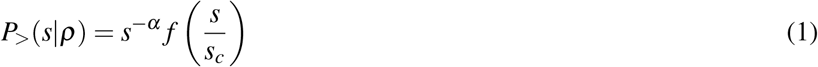

*ρ* is the protein concentration, *s* is the droplet size (either radius, diameter or volume) and *s*_*c*_ characteristic droplet size, defined as the ratio of the second to first moment of the size distribution, depends on *ρ*, and it is expected to diverge at the *ρ*_*c*_ critical concentration for phase separation as^43^

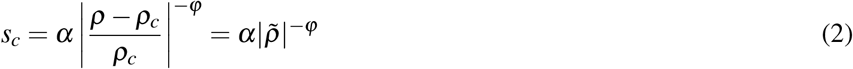

where *α*≥0 and *ϕ >* 0 are critical exponent, *a* is a constant and *f* is the scaling function. The analysis of experimental data by different research groups consistently resulted in critical exponents *α* = 0 and *ϕ* = 1, with the scaling function given by

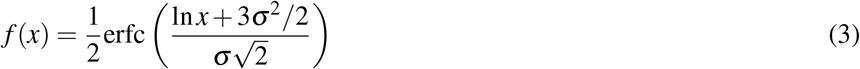

corresponding to a log-normal distribution of droplet size with standard deviation *σ*, independent of the density *ρ*. We stress that the exponents reported for standard percolation are *α* = 1.5 and *ϕ* = 2^44^. The deviation between the critical exponents suggests that proteins forming clusters below the critical concentration for phase separation may belong to a different universality class than standard percolation. Furthermore, the results are consistent with critical phenomena and continuous phase transition, consistently with earlier experimental results^45^.

Here we analyze whether fuzzy interactions enable a continuous transition from the dynamic clusters formed at sub-critical concentration to macroscopic condensates (Figure 2). The mathematical formalism for fuzzy interactions are detailed in Appendix A^46^. In the main text, we only discuss the key features of the model, which lead to the formation of condensates. Most importantly, a fuzzy interaction motif (Figure 1) can interact with various other motifs (*x*1, *x*2,…, *x*_*N*_) simultaneously with different membership functions (*m*(*x*_1_), *m*(*x*_2_), …, *m*(*x*_*N*_)) (Figure 1) within the framework of the fuzzy set theory^47^. An aromatic ring, for example, can interact both in parallel and perpendicular orientations with other *π* groups^48^. This leads to a set of simultaneous, partial contacts, which mutually influence the pairwise binding affinities (Appendix B). The likelihood of contacting all these potential interaction motifs depends on the length and dynamical properties of the segments in between the motifs^49^. In case the linker is fully dynamic, all potential interactions can be sampled, while a drop in dynamics also restrains the available interaction space. This is a key point, which is capable of describing a gradual change in binding affinity as the cluster grows^49^. Furthermore, considering the dissociation probability of the motifs, and the diffusion of the particles (monomers, oligomers, or large cluster/polymers) out of the interaction zone, the affinity can be tuned on-the-fly following the changes in occupancy (also considering partial sites). Simulations based this framework (Appendix A), correctly recapitulate that the fraction of oligomers and large polymers converge to a limit, instead of and infinite growth^49^ (Figure 2). The simulations also describe the assembly/disassembly of the clusters as well as their growth in size as the system approaches the phase boundary. Linker dynamics, through its impact on the number of available interaction sites, influence the rate, but not the extent of clustering (i.e. fraction of clusters)^49^ (Figure 2).

**FIG. 2.**
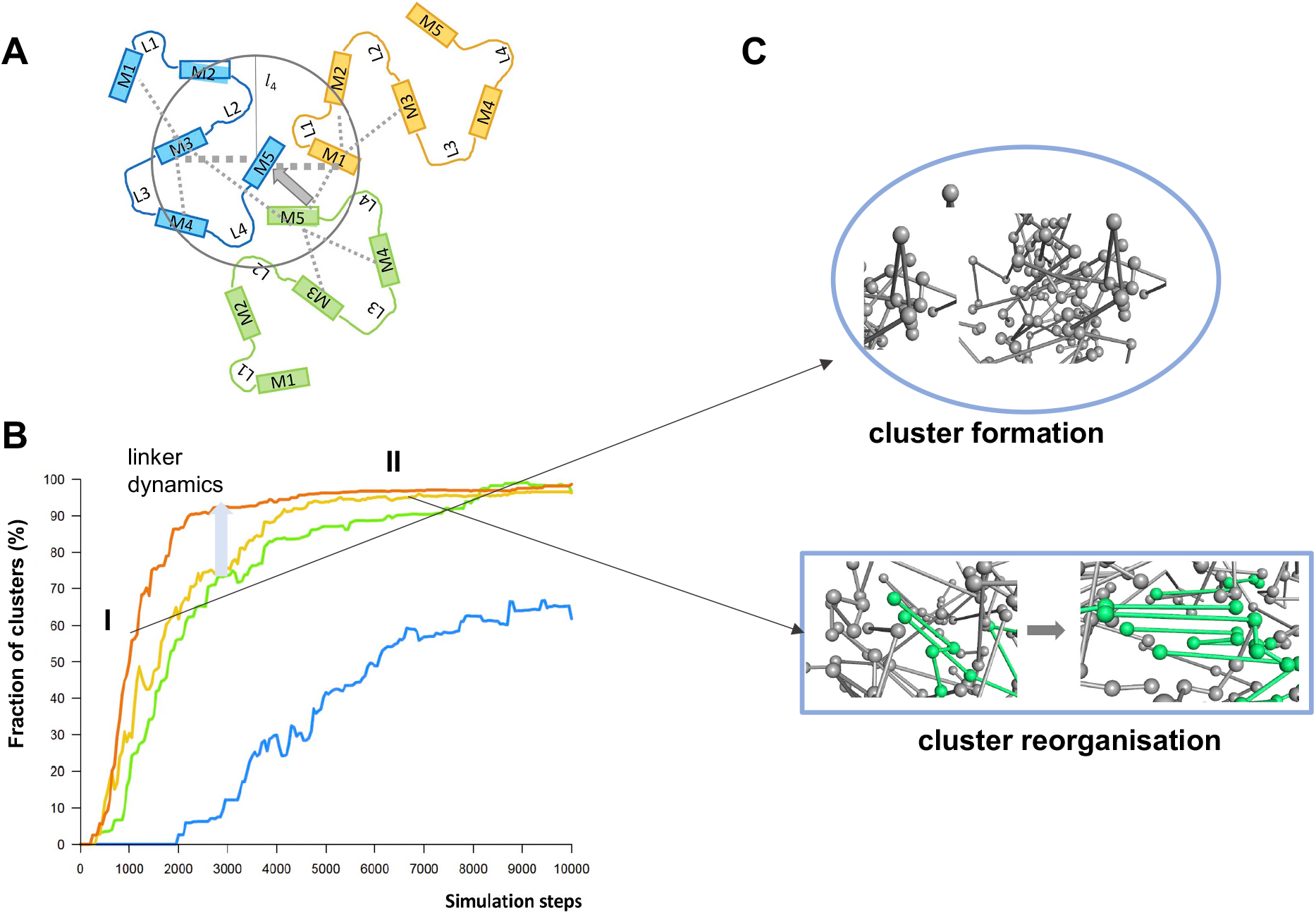
Simulating continuous phase transitions using a fuzzy interaction model. **A** The interactions are described using the fuzzy set theory, where partial contacts with multiple sites can be established simultaneously with different memberships. Different colors correspond to different monomers, where the putative binding motifs are marked by boxes (labelled M). Interactions with a given sphere are considered (circle), the size of which is determined by the dynamics of the linkers (marked by L). **B** Simulation trajectories of systems with different linker dynamics (blue low, red high, in arbitrary units). Cluster formation is considerably faster with increased dynamics. The simulations can be divided into two steps, cluster formation and growth (I), when both the number and size increases although the clusters dynamically assemble/disassemble; and reogranisation stage (II), when the fraction of clusters converged, but their internal structure continuously reorganise. In stage II the clusters still exchange monomers with the invironment. Simulation steps are given in arbitraty units. **C** Representative images from the two stages of the trajectory. During reorganisation, the contact pattern may adopt regular form.

In this section we conclude that a model based on fuzzy interactions (Appendix A) can describe continuous and dynamic cluster formation, converging to a phase separated system.

## IV. INFLUENCE OF THE ENVIRONMENT ON COLLECTIVE INTERACTIONS

Condensates are often formed/dissolved upon cellular cues or changes in solution conditions^16,50,51^. Their cellular context sensitivity also induces changes in the biophysical properties, for example in the material state of condensates^52^. This is of particular importance in case the biophysical properties perturb the physiological state^53^. Is the model based on fuzzy interactions can recapitulate the context-dependent behaviour of condensates? We follow two approaches, first based on the energy landscape framework adapted for protein interactions, and second based on computer simulations using the fuzzy model.

In analogy with the energy landscape framework for protein folding, the energy landscape of protein interactions can also be considered as rugged, instead of sampling only the distinguished *native* state^31^. Thus, proteins may sample alternative bound states while bound with their specific partners, which may be considered as non-native states. Each bound conformations are energetically sub-optimal, yet distinct with different proteins^54^, consistently with the notion of fuzzy interactions. Given this model, the bound state is represented by an ensemble of conformations and contact patterns, the populations of which depend on external factors. While variations in conditions shift the ensemble and may favor given bound forms, the bound state never confines to a unique form, but remains to be heterogeneous^31^. Therefore, the interaction between specific partners will respond to the conditions through changing the populations in the bound state ensemble^55^. Such change allows fine-tuning of biological activity, for example regarding substrate selectivity^54^.

Now we consider how the network of fuzzy interactions responds to external conditions. In the second part of the simulations based on the fuzzy interaction framework^49^ (Appendix A), which represents a phase separated system, monomers are still exchanging with the environment. The fraction of clusters remains stable, while their internal organisation may undergo substantial changes through variations in occupancies of the interaction sites, considering also partial contacts (Appendix A). The heterogeneous interaction network occasionally start forming regular interaction patterns, such as ladders of contacts formed by consecutive residues^49^. Depending on the linker dynamics, the network may transform from disordered to ordered interactions (Figure 2). Small perturbations of the model result in distinct outcomes of the simulations, illustrating the sensitivity of the system. Intriguingly, dynamic systems without fuzzy interactions do not exhibit such extensive rearrangements of the interaction network. We also note that context-dependent controls are often achieved through fuzzy inference systems^56^.

In this section we conclude that the network of fuzzy interactions is sensitive to the perturbations by external factors, thus the model may serve as a basis of collective, context-dependent behaviors.

## V. CORRELATION LENGTH AND CONTEXT-DEPENDENT BEHAVIORS

When a system develops a large correlation length − defined as the spatial range over which density fluctuations are strongly correlated −it behaves similarly to a system near criticality, at least on spatial scales smaller than the correlation length^20,57^. In the case of our droplet system, this correlation length could approach the size of individual cells as the critical density of the protein is approached from below. From statistical mechanics, it is well known that systems near criticality exhibit extreme sensitivity to external perturbations^58^. This means that even small changes can elicit prompt and widespread responses across both large spatial and temporal scales. In such a regime, fluctuations are not isolated; instead, they propagate rapidly through the system, allowing it to respond in a highly coordinated manner to external forces.

In biomolecular condensates, the situation is further complicated by the heterogeneity of the amino acids and their intra- and intermolecular microenvironments^59^. This suggests that if the system is indeed close to criticality − or equivalently, if it has developed a large correlation length −this molecular diversity might enable different parts of the system to respond distinctly and efficiently to various external stimuli. This may lead to distinct responses to different external signals, depending on the local conditions or the specific interactions between different protein components.

Within the framework of percolation theory, it is important to note that in standard percolation models, clusters form in an irreversible manner. Once a cluster forms, it remains intact, with no mechanism for the system to return to a prior state or for clusters to disintegrate. This lack of dynamical equilibrium distinguishes standard percolation from other processes where continuous formation and dissolution of structures occur. The scaling analysis suggests that the droplet formation observed in our system does not correspond to the behavior of standard percolation^19^. Instead, the dynamics we are studying seem to involve a more complex, reversible process. A more accurate and realistic model of this system would need to account for the ability of clusters or droplets to not only form but also to disintegrate and re-form dynamically.

This indicates that the system likely exists in a state of dynamical equilibrium, where clusters of proteins or molecules can fluctuate between different states in response to varying conditions, such as concentration, temperature, or other external factors. Thus, unlike in traditional percolation where the formation of large, connected clusters marks a one-way transition, the system we are dealing with appears to have more fluidity, with clusters constantly forming and dissolving. This suggests that models incorporating reversible aggregation and disaggregation would provide a more accurate description of the droplet formation process in our system.

## CONCLUSIONS AND OUTLOOK

In this article we argued that similar interaction mechanisms take place at various scales of protein assembly from oligomers to large clusters. We demonstrated that heterogeneous, variable interactions of specific partners may generate a collective network in a manner, which is consistent with continuous phase transitions. This in line with the scale-invariant log-normal distributions of the droplet sizes, which hints on second-order phase transition. The recently found validity of scale invariance place proteins close to their phase boundary, thus maximizing the sensitivity to environmental conditions. This is in accord with the model we presented here, which can describe the conversion between different material states. The importance of condensates in human physiology is increasingly recognized^9^. Likewise, increasing number of diseases are associated with perturbations of the biophysical properties of the droplet state^6,60^. In particular protein aggregation, a hallmark of various neurological disorders were related to the material state conversion of liquid-like condensates^61^. The premise is that understanding the driving forces and the nature of collective interactions may open novel approaches for target selection or drug development for condensation diseases. The results indicating the proximity of proteins close to their phase separation boundary also implies that the fluctuations in the system can be efficiently modulated, also by small-molecule drugs. Promising attempts towards this direction have been published^62^. Another possible application is to exploit context-sensitivity for modulating cellular pathways^63^, which eventually can be used as a readout in screening studies. While there is a long, rugged road ahead (such as the interaction energy landscape of the condensates), quantitative models may contribute to all these efforts.

## ACKNOWLEDGEMENT

This work was supported by AIRC IG 26229 (M.F.) and PRIN 2022EMZJL4 (M.F.).

## APPENDIX A

This appendix describes a mathematical formalism for fuzzy binding and its application to formation of higher-order assembly. More details can be found in the reference^49^.

### Definition

#### Binding of two molecules

Two molecules were associated if at least 1 interaction was established between two motifs, which were located on separate molecules.

#### Oligomers

Oligomers contained (1 *< n <* 25) molecules, each of which was connected to the rest of the cluster via at least 1 motif-interaction.

#### Polymers

Polymers contained *n* ≥25 molecules, each of which was connected to the rest of the cluster via at least 1 motif-interaction.

#### Aggregates/fibrils

Amyloid-like elements were defined based on a regular interaction pattern amongst *n* 5 ≥molecules. Regular interaction patterns were connections between at least 2 consecutive binding elements on both interacting partners.

### Theoretical framework

#### Interaction between two motifs

Binding affinity between two motifs was characterized by the average of the individual binding preferences:

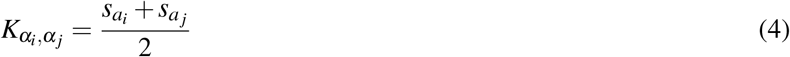

where 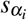and 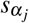are the binding preferences of the interacting *α*_*i*_ and *α*_*j*_ motifs. Two motifs where bound if 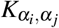above a given threshold.

#### Potential interactions with multivalent sites

In the fuzzy framework, partial interactions with multiple sites are also considered. First, we define the volume, within which potential binding elements were located:

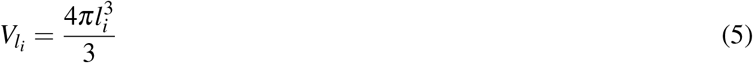

where *l*_*i*_ is the length of the linker, and the spherical volume 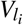 is centered on the binding element *α*_*i*_. Then the number of binding elements, which were located within this volume was determined:

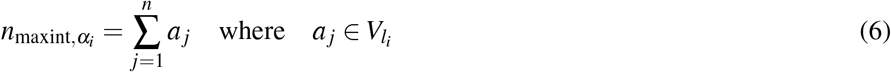

where 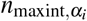 is the number of binding motifs, which have at least 1 residue within the defined volume. 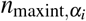 is the maximum number of the potential interaction sites, which are available for interactions with *α*_*i*_ motif. The number potential binding sites also modulate the interaction affinity between two motifs (*α*_*i*_, *α*_*j*_).

#### First association event (anchoring step)

The association probability in the first step is defined as:

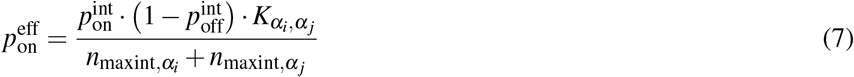

where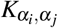 is the binding affinity between *α*_*i*_ and *α*_*j*_ motifs (eq. 4), 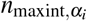 and 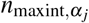are the maximum number of potential interaction elements around *α*_*i*_ and *α*_*j*_ motif, respectively. Competing motifs reduce the association probability between two motifs (*α*_*i*_, *α*_*j*_).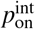 and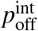are the intrinsic association and dissociation probability. The binding is realized 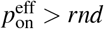 where *rnd* is a random number.

#### Determination of occupancy

After the first binding step, an occupancy of each binding element is determined. As the applied framework also considers partial interactions, a fuzzy union (s-norm) operator (based on a fuzzy set theory^47^) for each connected motif is determined:

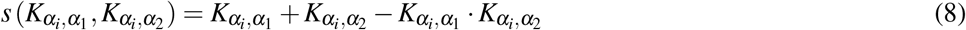

where 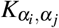is the binding affinity of *α*_*i*_ and *α*_*j*_ motifs (eq. 4). The occupancy of each motif is defined as and algebraic sum:

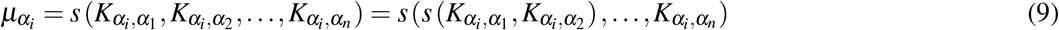

where 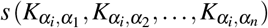 is the fuzzy union operator in eq. 8. Occupancy (eq. 9) is computed for each, even partially bound motifs (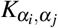 above a threshold).

### Subsequent binding events

#### Modified binding affinity

Once two motifs were attached, the binding affinity between two motifs (eq. 4) is modified by the corresponding occupancies of potential binding sites:

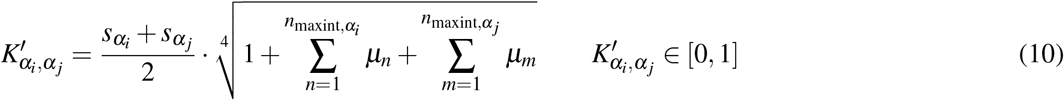

where *µ*_*n*_ and *µ*_*m*_ are the occupancies of all potential binding elements (eq. 6) within the given volume (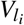) around *α*_*i*_ and *α*_*j*_ (eq. 5). Thus, high occupancies of the potential interaction motifs within 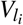 reduce the competition for *α*_*i*_ and *α*_*j*_, respectively, and concomitantly increase 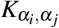 affinity.

#### Modified association probability

After the first binding event (once the two motifs were bound) the association probability was also modified by the corresponding occupancies:

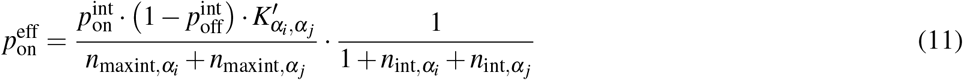

where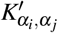 is the modified binding affinity defined in eq. 10, 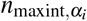 and 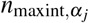 are the actual number of binding elements, which are bound to *α*_*i*_ and *α*_*j*_, respectively.

#### Dissociation

The dissociation probability is defined as (inverse relationship to the binding affinity):

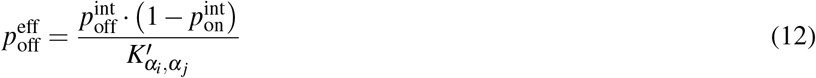

where the intrinsic association 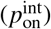 and dissociation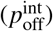probabilities are the same as in eq. 7, and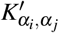 is the modified binding affinity defined in eq. 10.

#### Diffusion

The different kinds of species (individual molecules, oligomers and polymers) can be transformed into one of the neighboring boxes with a probability:

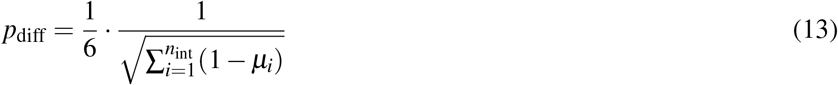

where (1 − *µ*_*i*_) is the available interaction capacity of a given *α*_*i*_, binding element. Summation is carried out for all motifs, which are bound to 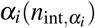. Therefore, larger polymers with more available binding capacity are less likely to move to another box. For simplicity, during diffusion, the molecules/oligomers/polymers are assumed to move as rigid units.

#### Linker dynamics in fuzzy models

Within the framework of the fuzzy interaction model, we describe multivalent/multisite binding trough potential contacts with interaction sites (eq. 6) within the 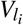 volume (eq. 5). This volume is dependent on the linker length and dynamics. Both the association probability (eq. 11) and binding affinity (eq. 10) between two motifs were dependent on 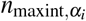 (eq. 6), thus are indirectly influenced by the linker properties. If the linker is infinitely flexible the neighboring motifs can sample the whole volume (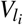, eq. 5), which is available for interactions. Less dynamical linkers, however, cannot visit the whole interaction space, which can be expressed through a modified linker length:

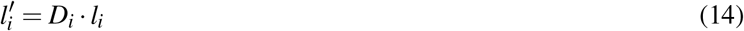

where *D*_*i*_ is the linker dynamics and *l*_*i*_ is the length of the linker.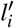is used to obtain the volume available for interactions (eq. 5).

### Computational protocol for simulations using the fuzzy model

In the first simulation step, binding between multivalent motifs could take place according the association probabilities (eq. 7). Once the initial contacts are made, the occupancies are determined in each step. The binding affinities (eq. 10) and association probabilities (eq. 11) are modified accordingly. In all subsequent steps, binding or dissociation of an interaction element from the other available or bound motifs can be decided based on the association (eq. 11) and dissociation probabilities (eq. 12). Each species (individual molecules, oligomers and polymers) can be moved to the one of the neighboring boxes according to the diffusion probabilities (eq. 13). The simulation steps are given in arbitrary units. Using a contour length of 4Å/residue^64^, and an approximate diffusion constant^65^ *D*∼ 1.1× 10^−6^cm^2^s^−1^, a approximate simulation step corresponds to a *µ*s motion (rough estimate for individual molecules).

## APPENDIX B

This appendix describes a mathematical formalism for determining the universal coefficients of the scale invariant model (eqs.1,2 main text). More details can be found in the reference^19^.

To determine the exponents *α* and *ϕ*, we introduce the moments of *P*_*>*_(*s*|*ρ*) for *k >* 0

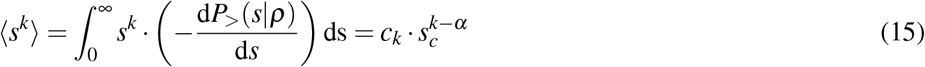

where the scaling ansatz (eq. 1) has been used in the last step and *c*_*k*_ is given by

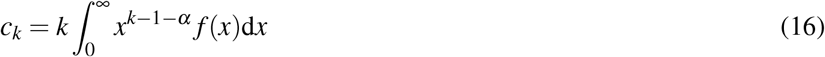

which depends on the function *f* but is independent of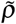. From eq. 15, we deduce that

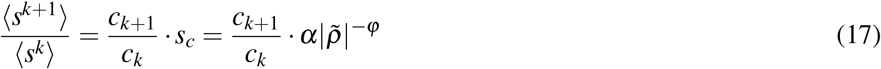

If the scaling ansatz of eq. 1 is correct, by plotting the ration of moments ⟨*s*^*k*+1^⟩ */* ⟨*s*^*k*^⟩ for various values of *k* as a function of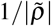in a double logarithmic scale, we should obtain straight parallel lines with slope *ϕ* and intercept ln(*ac*_*k*+1_*/c*_*k*_). Having determined the exponent *ϕ*, we can also determine the exponent *α* using eq.15

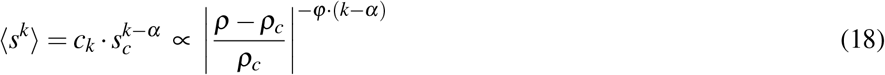

The exponent *α* is the calculated from

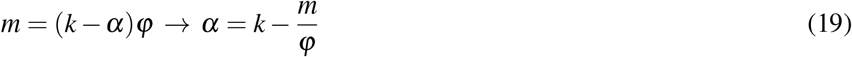

where *m* is the slope of the linear fit of the double logarithmic plot of ⟨*s*⟩ (the first moment, *k* = 1) vs 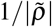. We then plotted the mean droplet size, ⟨*s*⟩, versus the inverse distance from the critical concentration 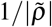 on a natural logarithm scale.

Scale invariance could be probed in two steps: *i)* first determining the average (*s*_0_) and variance (*σ*) of the log-normal distribution, *ii)* second, probing that the variance of the log-normal distribution, *σ* is independent of the concentration *ρ*. The scaling ansatz of eq. 1 for the SDF is equivalent to the following scaling for the probability density distribution

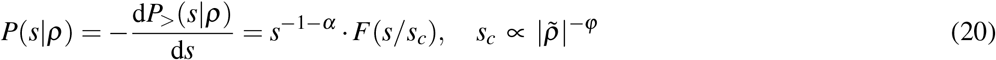

where *f*, the scaling function in eq. 1, and *F* are related as follows

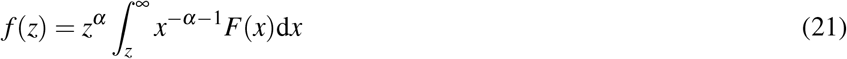

The log-normal droplet size distribution *P*(*s*|*ρ*) is

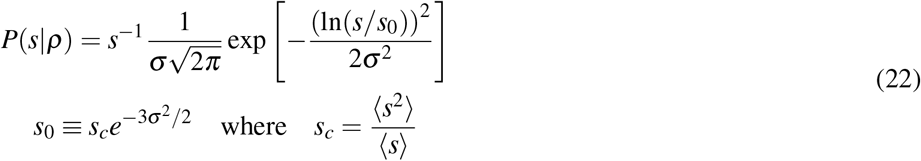

being the characteristic droplet size as defined above. Consequently, the size survival distribution function is

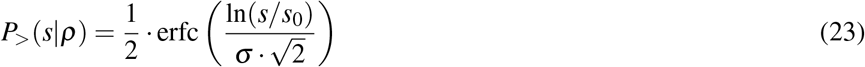

The values of *s*_0_ and *σ* can be determined as the average and the variance, of ln(*s/u*) obtained at each concentration:

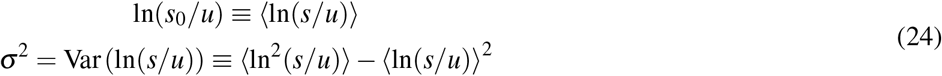

where *u* is an arbitrary (and irrelevant) constant with the same units as *s*. In the following, when not stated, it is implicitly assumed that *u* = 1 in the same units as *s*. The droplet sizes follow a log-normal distribution only if the survival distribution functions or, equivalently, the size distribution functions, multiplied by *s*, collapse when plotted versus eq. 22 or eq. 23 with the values of *s*_0_ and *σ* of each droplet size distributions obtained at different concentrations (eq. 24).

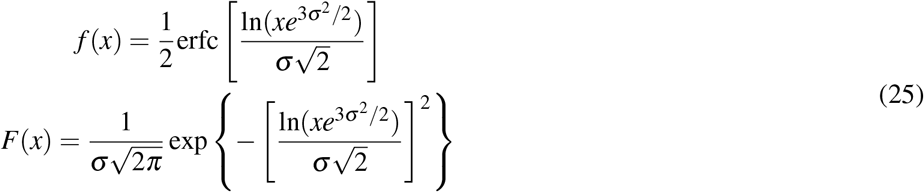

with *α* = 0 and *ϕ* = 1. Notice that the log-normal distribution implies that *α* = 0 whereas the value of *ϕ* is not determined a priori. Furthermore, the *k*−th moment of the log-normal distribution is

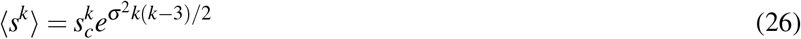

which, compared with the scaling prediction eq. 15, is also consistent with *α* = 0, appropriate for the log-normal, and

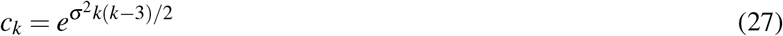

which is independent of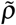 if *σ* is independent of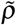, see eq. 15.

